# African Pan Genome Contigs Expose Biologically Relevant Sequence Still Hidden from Human Reference Frameworks

**DOI:** 10.1101/2025.08.15.670543

**Authors:** Rachel Martini, Abdulfatai Tijjani, Kyriaki Founta, Daniel Cha, Alexandria Awai, Sebastian Maurice, Jason A. White, Christopher E. Mason, Isidro Cortes-Ciriano, Nicolas Robine, Onyinye Balogun, Nyasha Chambwe, Melissa B. Davis

## Abstract

Human reference genomes underpin biomedical discovery but remain incomplete and biased toward European populations, constraining interpretation of genetic variation in underrepresented populations. Here we characterize African Pan Genome (APG) contigs totaling 296.5 Mb to define the sequence and functional landscape of genomic regions absent from current references. Most contigs align to the telomere-to-telomere (T2T-CHM13) genome and across 47 haplotype-resolved Human Pangenome Reference Consortium (HPRC) assemblies, with T2T-CHM13 placements enriched in centromeric and satellite repeats and overlapping 373 genes, including disease-associated loci. Mapping across HPRC assemblies revealed ancestry-associated contig enrichment, particularly in African genomes. Notably, 742 contigs remained unmapped under both stringent and relaxed criteria. These sequences are largely nonrepetitive and exhibit strong functional potential, including predicted protein-coding genes, CpG islands and transcriptional activity. Together, these results demonstrate that functionally relevant, ancestry-enriched genomic sequences remain absent from current references, with important implications for disease variant interpretation and precision medicine.

## Introduction

The human reference genome determines which sequences align, what variants are called and annotated, and ultimately what biological data becomes visible to genomic research studies. Despite the availability of newer reference genome assemblies, GRCh37 and GRCh38 remain the primary references used for sequence alignment in human genomics research and clinical practice^1,2^. With unresolved gaps comprising approximately 7% of the GRCh38 genomes^3,4^, it reflects a limited view of human diversity, primarily focusing on European ancestry^5^. This Eurocentric reference bias affects genomic medicine by essentially rendering data that is relevant to specific ancestry groups invisible^1,2,6,7^. A clear illustration of this bias comes from genome-wide association studies (GWAS) that use platforms designed from a Eurocentric reference, have historically been conducted in primarily European populations, and lead to limited applicable replication across any other ancestries^8–11^. This underrepresentation can hinder the mapping of genetic risk alleles, our understanding of regulatory gene networks that define disease biology and therefore limit the precision of genomic medicine.

Advances in long-read sequencing are transformative for genome science by overcoming the limitations of existing human reference genomes, resolving previously inaccessible regions of the genome. The Telomere-to-Telomere (T2T-CHM13) assembly provides a complete, gapless linear reference genome, including centromeres and segmental duplications^3^. However, both GRCh38 and T2T-CHM13 remain limited in ancestral diversity^5,12^. More recently, the Human Pangenome Reference Consortium (HPRC) has introduced a more diverse reference framework, initially generating 47 assemblies with wider representation across genetic ancestry groups. These assemblies also capture population structural variation allowing the building of a pangenome graph that offers better sensitivity in variant calling across diverse populations^4,13^.

The African Pan Genome (APG) study by Sherman et al^14^ provided a powerful test case for this challenge by identifying 296.5 million base pairs missing from GRCh38, prior to the release of the T2T-CHM13 and HPRC reference genomes. This landmark study performed whole-genome sequencing of 910 individuals of African descent, and subsequently assembled unmapped GRCh38 reads into 125,715 contigs^14^. While these contigs revealed previously uncharacterized genomic content specific to individuals of African ancestry, their biological importance and overlap with newer reference builds remained unclear.

In this study, we investigated whether unresolved APG contig sequences, absent in GRCh38, reflected low-complexity sequence of limited consequence or biologically meaningful genomic content still excluded from modern reference frameworks. We asked three questions: (1) how much ancestry-enriched sequence absent from GRCh38 is recovered by T2T-CHM13 and HPRC assemblies; (2) whether recovered sequence overlaps functional genomic elements; and (3) whether contigs still missing from current references show evidence of coding, regulatory, or transcriptional activity. We determined that the majority of these contigs correspond to novel, functionally relevant regions of the genome. By evaluating the representation and functional roles of these sequences, our work emphasizes the risks of missing essential data in genomic research when we continue to rely on incomplete and Eurocentric reference genomes. Despite the options of more comprehensive reference genomes, GRCh38 is still the predominant genome reference used to date, which leads to persistent blind spots in our understanding of genome function, and perpetuates the omission of ancestry-specific biologically and clinically significant genomic content.

## Results

To determine how much ancestry-enriched sequence remains outside of the current reference genome frameworks, we aligned 124,240 contigs assembled by Sherman et. al^14^ against T2T-CHM13 and 47 HPRC linear assemblies and then layered functional annotation, repeat analysis, and RNA-seq evidence onto mapped and unmapped subsets **(see Methods, Figure 1, Figure S1)**. This design allowed us to distinguish sequences recovered by modern references from sequences that remain unresolved even after gapless reference genome advances.

**Figure 1.**
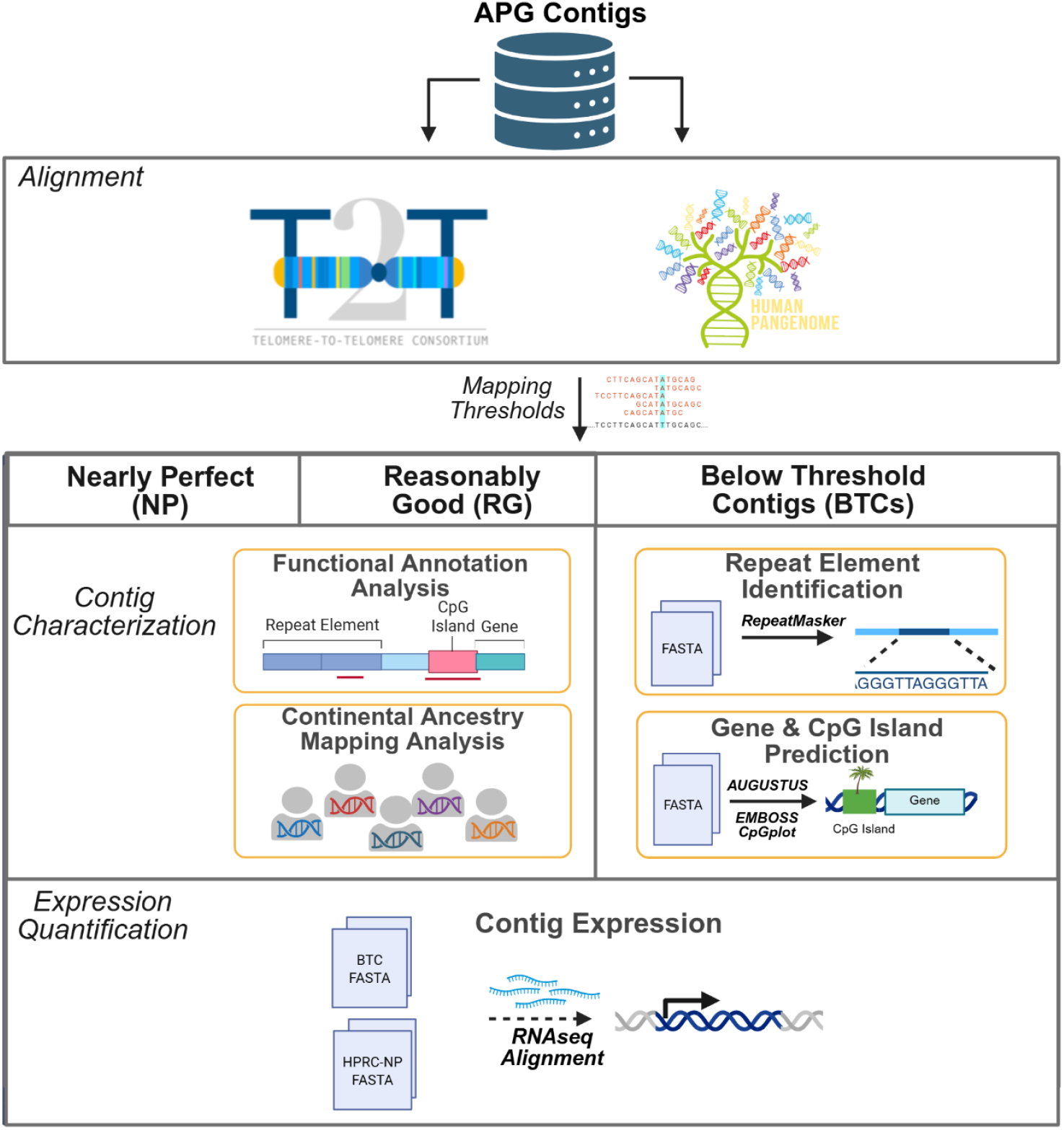
Study Design. APG contigs were aligned to the T2T-CHM13 reference genome and to 47 haploid paternal linear assemblies from the Human Pangenome Reference Consortium (HPRC; Year 1, Release 1). Based on predefined mapping quality thresholds, alignments were classified as “Reasonably Good” or “Nearly Perfect”. Contigs that failed to align, or that produced only weak alignments to both T2T-CHM13 and HPRC references, were designated as “Below Threshold Contigs (BTC)”. Contigs with “Nearly Perfect” alignments to T2T-CHM13 were further examined based on their functional annotations, whereas those aligning “Nearly Perfectly” to HPRC assemblies were used for continental ancestry analysis. For “Below Threshold Contigs (BTC)”, we performed de novo gene prediction and CpG island identification, as well as quantification of expression of predicted genes. Repeat element analysis was conducted on both the “Below Threshold Contigs (BTC)” and the “Reasonably Good” alignments to any of the reference genomes.

### APG contigs map to T2T-CHM13 regions absent from GRCh38

We first determined whether the 124,240 APG contig sequences missing from GRCh38 were recovered in the gapless T2T-CHM13 reference assembly. We found that 39.50% (49,070/124,240) of the APGs aligned to T2T-CHM13 at the Nearly Perfect (NP) threshold, with 94.45% of these T2T-CHM13 alignments mapping to regions unique from GRCh38 **(Figure 2A-B, Tables S1-3)**. Using the more relaxed Reasonably Good (RG) threshold increases the recovery of APG contigs mapping by 47.77% and likely reflects complexity in variation not represented as much in the assemblies available. The highest number of alignments to T2T-CHM13 were in Chromosomes 9, 21, and 14, and the least number of alignments to Chromosomes 18, 16 and Y **(Figure 2C)**. As anticipated, recovery was driven largely by previously unplaced contigs, where neither end of the contig sequence was previously resolved to a location on GRCh38^14^ **(Figure 2D)**, underscoring how unmappable sequence to legacy references misses real sequence.

**Figure 2.**
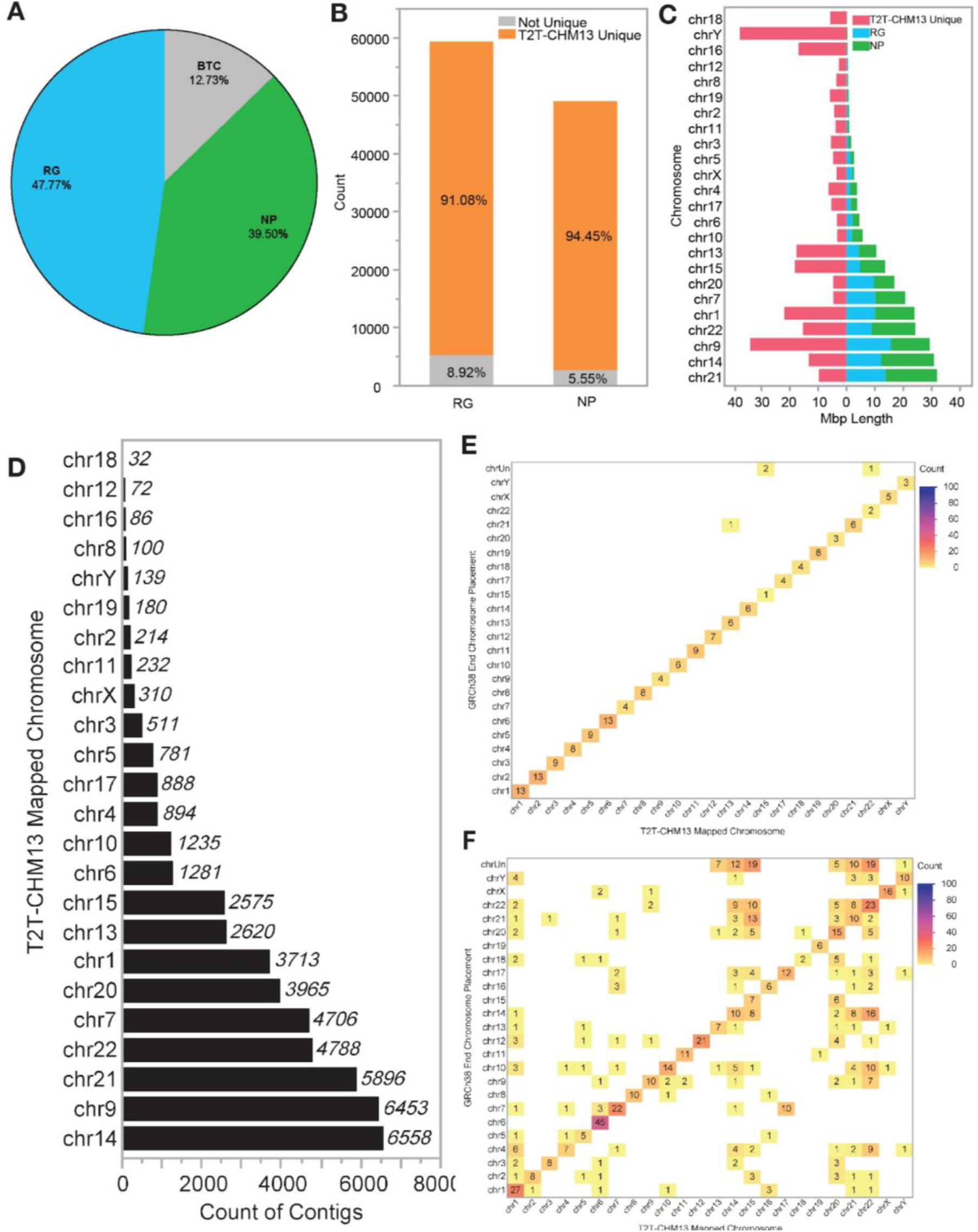
Summary of APG Contig Alignment to T2T-CHM13. (A) Pie chart showing distribution of contigs with Nearly Perfect (NP, green), Reasonably Good (RG, blue) or below threshold (BTC, gray) alignments to T2T-CHM13. (B) Stacked bar chart depicting T2T-CHM13 RG or NP alignments that map to T2T-CHM13 unique regions compared to the GRCh38 reference (orange). Alignments to the non-unique areas are shown in gray. (C) Butterfly plot of T2T-CHM13 alignment counts per chromosome (right) and total aligned length (Mb) T2T-CHM13 unique content from GRCh38 (left, pink). All alignments shown passed either the RG (blue) or NP (green) thresholds. (D) Bar chart of counts of unplaced APG contigs mapping to T2T-CHM13 chromosomes. Heatmaps of GRCh38 chromosome placements and T2T-CHM13 chromosome NP mapping results for (E) Two-End placed and (F) One-End placed APG contigs.

### Re-evaluating APG contig placement from GRCh38 using the T2T-CHM13 reference

We further assessed APG contig alignment to the T2T-CHM13 assembly in the context of their prior placement status on the GRCh38 reference genome **(Table S4)**^14^. Of 124,240 contigs, 302 were previously classified as two-end (TE) placed, 1,246 as one-end (OE) placed, and 122,692 remained unplaced. Unplaced contigs showed the highest level of reasonably good (RG) alignments (48.08%), yet the lowest proportion of NP alignment (39.1%). A lower proportion of OE (26.00%) and TE (12.25%) had RG alignments, but had similar substantial NP alignment (51.32% and 55.06%, respectively) **(Table S4)**. These observations indicate that the contigs with greater sequence divergence were harder to align.

Chromosome concordance was nearly complete for TE contigs, with only one discordant NP alignment **(Figure 2E)**. Three TE placed contigs that had been placed on GRCh38 unknown chromosomes mapped to T2T-CHM13 Chromosomes 15 and 22 (n = 2, n = 1, respectively). In contrast, 53.21% of OE contigs with NP alignments mapped to different chromosomes in T2T-CHM13 than in GRCh38 **(Figure 2F)**, and 64.89% of these discordances involved acrocentric chromosomes. Similar patterns of discordant alignments were also observed among RG mapping contigs **(Figure S2)**. In line with our expectations, we find that the vast majority of APG contigs map to the T2T-CHM13 genome at locations that correspond to novel sequences not present in earlier versions of the reference genome.

### APG contigs are recovered in gaps resolved by T2T-CHM13 that are repeat-rich and functionally important

As the overwhelming majority of the NP contigs (94.45%) mapped to regions unique to T2T-CMH13 **(Figure 2B)**, we sought to determine their potential overlap with known functional annotations. Most APG contigs recovered in T2T-CHM13 overlap centromeric and satellite-rich sequence (94.16%), consistent with longstanding limitations of short-read-based reference construction. The remaining contigs were annotated to genes (2.64%), CpG islands (3.61%) and composite elements (4.39%) **(Figure 3A, Table S5)**, with similar enrichment patterns for the less stringent RG-aligned contigs **(Table S5, Figure S3)**.

**Figure 3.**
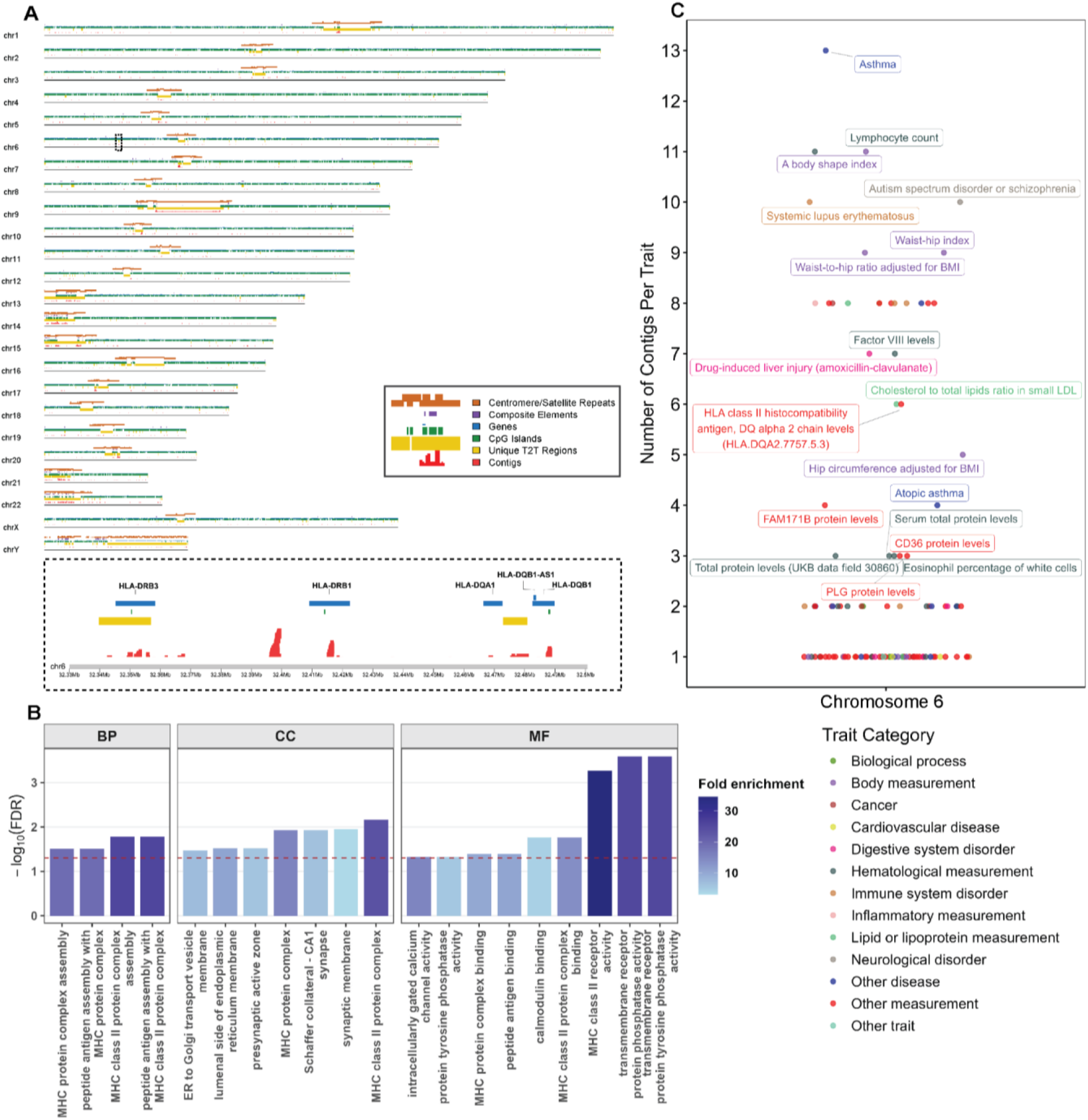
Distribution of Functional Feature Overlap of T2T-CHM13 Mapped Contigs. (A) Placement of nearly perfect aligned contigs across T2T-CMH13 chromosomes overlaid across various annotation categories. Contigs are highlighted in red. Enrichment was examined across T2T-CMH13 unique regions in comparison to GRCh38 (yellow), CpG Islands (green), RefSeq annotated genes (blue), composite elements (pink), and centromeres/satellite repeats (brown). A zoomed-in 170 Kb region (black dashed-line box) at the chromosome 6 HLA locus with the highest number of mapped contigs across genes is shown at the bottom of the panel. (B) Significantly overrepresented Gene Ontology terms (BP: Biological Processes, CC: Cellular Component, MF: Molecular Function) among the 373 T2T-CMH13 genes with at least one APG contig nearly perfectly aligned to them. Bars represent enrichment significance as - log10(FDR), and color indicates fold enrichment. (C) Overlap of APG contigs with GWAS hits on chromosome 6. Each dot represents a GWAS trait, with the y-axis indicating the number of contigs overlapping with GWAS hits (polymorphisms) for that trait. Trait labels are shown, and all traits are colored according to their parent trait category.

T2T-CHM13-mapped APG contigs also overlap 373 annotated genes, with enrichment in immune, synaptic, and intracellular signaling pathways. Specifically, we found statistically significant enrichment in biological processes related to antigen presentation via the MHC class II protein complex. Cellular component and molecular function gene ontology processes related to neuronal recognition and the regulation or modulation of synaptic signaling were also enriched **(Figure 3B, Table S6)**. Interestingly, MHC class II enrichment was driven by the HLA locus on chromosome 6, where 84 contigs mapped to this 170Kb region on chromosome 6, with 44 contigs mapping to 4 HLA genes (*HLA-DRB3, HLA-DRB1, HLA-DQA1, HLA-DQRB1*) **(Figure 3A)**; in line with the known challenges of read mapping to this highly polymorphic region^15–17^. The main driver of contig overrepresentation in neuronal signaling processes was liprin-alpha 1 (*PPFIA1)*, a scaffold protein involved in synaptic organization^18^, with 33 contigs mapping.

We also determined contig-gene overlaps with disease phenotypes in the Online Mendelian Inheritance in Man (OMIM) database^19^ and GWAS-linked traits. Specifically, 234 contigs mapped to 60 distinct genes associated with OMIM disease phenotypes (**Table S7**), and 33 contigs overlapped with 113 previously identified GWAS hits (**Figure S4A, Table S8**). Contig–GWAS hit overlaps were detected across multiple chromosomes, with enrichment on chromosome 6, where 23 contigs overlapped GWAS hits linked to 23 distinct phenotypic traits (**Figure 3C)**. Asthma was the top enriched GWAS trait with 13 Contig-GWAS locus hits. This enrichment may reflect that the APG contigs were derived from analysis of an asthma cohort (Consortium on Asthma among African-ancestry Populations (CAAPA))^14,20^. Asthma was the second most enriched trait in RG mapped contigs, following autism spectrum disorder or schizophrenia (91 Contig-GWAS hits) (**Figure S4B**). These results imply that functionally important traits or disease-associated genetic variations in more diverse populations may be missed when using older versions of the human reference genome due to mapping challenges, particularly in the context of short-read datasets.

### HPRC assemblies recover APG contigs across diverse ancestral references

Next, we aligned the APG contigs to the 47 haploid paternal assemblies from the Human Pangenome Reference Consortium (HPRC) v1^4^, which capture a broader range of population diversity than single linear references, including complete genomes from African populations. Of the 47 HPRC assemblies, 24 are derived from individuals of African ancestry (AFR, **Table S9**).

Compared to T2T-CHM13 alignments, we observed a substantial improvement in APG contigs placement. Overall 99.40% of the APG contigs achieved a RG alignment, and 82.91% met the NP alignment threshold to at least one HPRC assembly (n = 123,498 and n = 103,003, respectively) **(Table S10)**. Notably, 80.44% of APG contigs have at least 1 NP alignment to an AFR ancestry HPRC assembly (n = 99,944) **(Table S10)**.

Across all HPRC linear assemblies, mapping improvements were observed across previously defined GRCh38 placement categories. At the NP threshold, 95.03% of TE contigs, 94.06% of OE contigs, and 82.76% of unplaced APG contigs aligned to at least one HPRC assembly (n = 287, n = 1172, n = 101,544, respectively) **(Table S11-S13)**.

### APG contig alignments to HPRC assemblies reveal population-associated enrichment

Comparing NP alignment results across reference assemblies, 49,020 contigs mapped to both T2T-CHM13 and the HPRC linear assemblies, whereas 53,983 contigs showed NP alignment exclusively to specific HPRC assemblies **(Figure 4A)**. Among these HPRC-only contigs, we observed a notable enrichment of alignments to assemblies from the AFR superpopulation (n = 7,969) (*p* < 2.2 e-16), whereas alignments to the other reference superpopulations were less frequent **(Figures 4B, Figure S5)**. Odds were 1.53x higher than to American (AMR) assemblies, and substantially greater versus East Asian (EAS, OR=2.74), European (EUR, OR=7.23), and South Asian (SAS, OR=8.55) assemblies **(Figure 4B)**.

**Figure 4.**
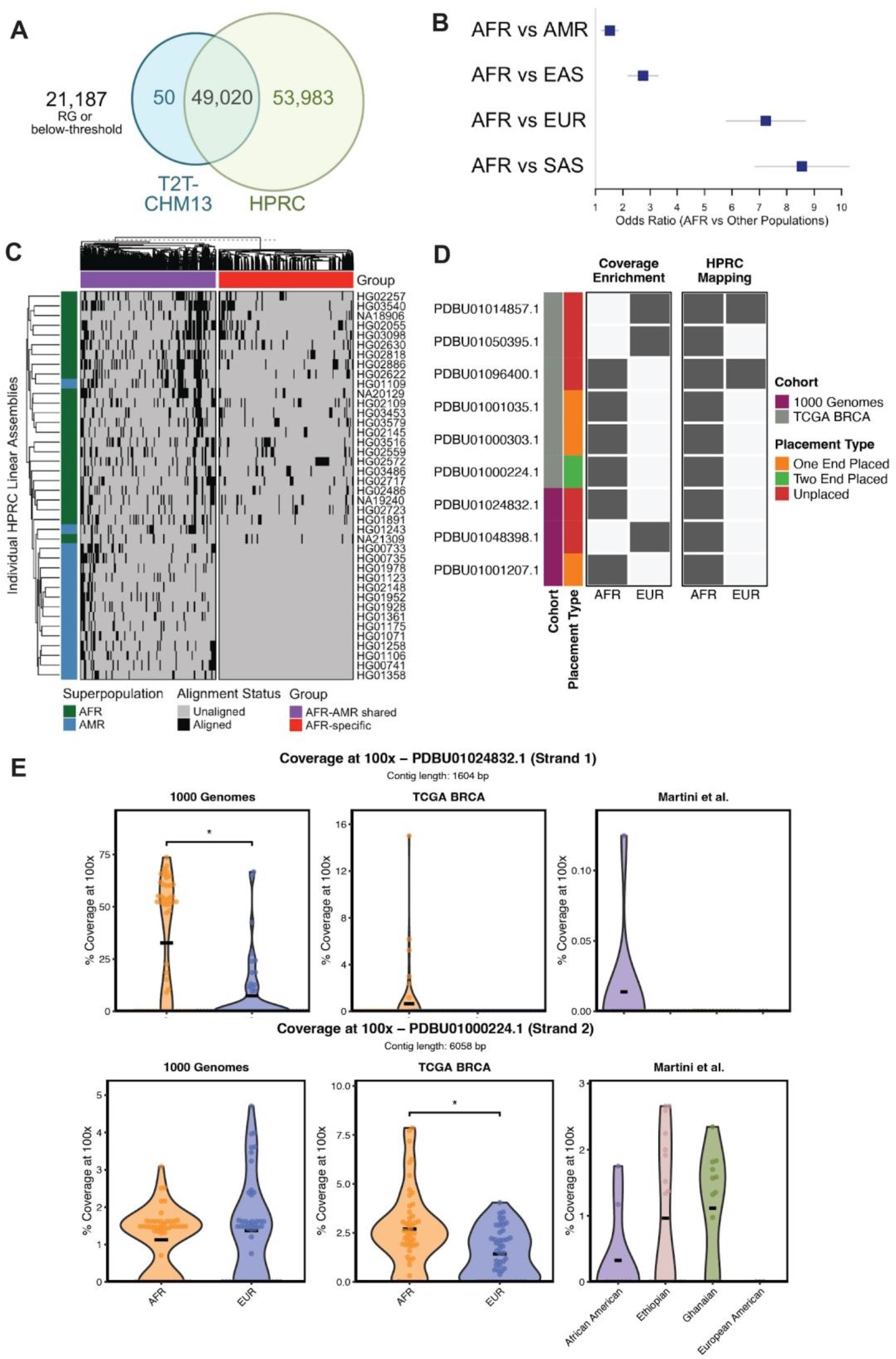
Summary of APG Contig Alignment and Expression Patterns across HPRC Assemblies. Venn diagram depicting the overlap of NP APG contig mapping to T2T-CHM13, HPRC linear assemblies. (B) Odds ratios (OR) comparing the likelihood of APG contigs alignment in African (AFR) genomes versus other superpopulations. (**C**) Heatmap showing alignment across individual linear assemblies from African (AFR) and Admixed American (AMR) HPRC samples. (D) Characterization of the 9 contigs with differential RNAseq read coverage between AFR and EUR samples across 1000 Genomes and TCGA BRCA. Each row represents a contig, while the left color bar indicates the cohort in which it was identified as significant (purple, 1000 Genomes; blue, TCGA BRCA). HPRC mapping information across AFR and EUR assemblies, as well as exclusive AFR assembly mapping status and placement type relative to GRCh38 are shown for each differentially enriched contig (dark grey indicates a positive hit). (E) Violin plots showing the percentage of contig length covered at ≥100× RNA-seq coverage depth across all 3 studied cohorts for 2 of the 9 contigs identified as differentially covered between AFR and EUR individuals: PDBU01024832.1 (identified in 1000 Genomes cohort; top) and PDBU01000224.1 (identified in TCGA BRCA cohort; bottom). Each point represents an individual sample, and samples are stratified by population group. Crossbar shows the mean of the distribution. Asterisks denote statistically significant differences between groups (*p-*value less than 0.05 (*), 0.01 (**) or 0.001 (***)).

Hierarchical clustering of the alignment patterns further revealed ancestry-associated grouping of individuals, with AFR assemblies clustering together and AMR assemblies forming a distinct secondary cluster of contig alignments **(Figure 4E)**. Unsupervised clustering of alignment patterns further highlighted contigs that mapped predominantly or exclusively to the AFR and AMR populations **(Figure S5A)**. The largest overlap between any two superpopulations consisted of 8,006 contigs that aligned to both the AFR and Admixed American (AMR) populations (**Figure S5B**). Although, AFR and AMR assemblies are more heavily represented in the HPRC v1 dataset (n = 24 and n = 16, respectively), this contig alignment overlap supports the population-associated nature of the APG contigs and suggests shared ancestry signals or structural variation between the AFR and AMR superpopulations. In contrast, shared mappings to the EAS, EUR, and SAS populations were considerably smaller, possibly indicating a limited representation of APG contigs within these genomes **(Figures 4B-C)**.

### Expression quantification of the HPRC-mapped only contigs

We quantified strand-specific transcriptional coverage of the 53,983 contigs that mapped exclusively to HPRC assemblies using RNAseq data from three independent cohorts: 100 samples respectively from the 1000 Genomes Project (1KGP)^21^ and the (TCGA) breast cancer (BRCA) project^22^, and 44 samples from an African ancestry-enriched breast cancer cohort (Martini et al.)^23^. The 1KGP and TCGA cohorts included approximately balanced African (AFR) and European (EUR) ancestry samples, whereas the Martini et al. cohort was enriched for African ancestry samples (13 Ghanaian, 19 Ethiopian, 9 African American, and 3 European American patients, see **Methods**).

Overall, only a small fraction of contigs exhibited detectable transcription. At the lowest read depth threshold (T1, ≥1 read), approximately 5% of contigs showed non-zero strand-specific coverage in at least one sample from 1KGP, with comparable rates in TCGA BRCA (**Figure S6A**). In the Martini et al. cohort, Ethiopian samples exhibited the highest detection rate (∼5.5%), followed by African American, Ghanaian, and European American samples. Detection rates decreased with increasing read depth stringency across all cohorts, with fewer than 1% of contigs retaining detectable coverage at T1000 or above (**Figure S6A**). These patterns were consistent across both the forward and reverse strands.

We next assessed ancestry-associated differences in contig coverage using strand-specific Wilcoxon rank-sum tests at the T100 threshold with Benjamini-Hochberg correction (**Figure S6B, Figure 4D**). In 1KGP, two contigs reached significance on the forward strand, including one enriched in AFR (PDBU01024832.1) and one in EUR (PDBU01048398.1), with additional AFR-enriched signals observed on the reverse strand (**Figure S7**). In TCGA BRCA samples, a larger number of differentially covered contigs were identified on the reverse strand, predominantly enriched in AFR (**Figure S5B, Figure S8**).

Among 9 contigs showing differential coverage between the two populations (**Figure 4D**), all mapping to AFR HPRC assemblies, whereas only 2 had previously been mapped to EUR assemblies. Notably, one contig (PDBU01000224.1) mapped exclusively to AFR HPRC assemblies and not to any other reference population assembly, consistent with its enriched expression in AFR samples, included Ethiopian and Ghanaian samples in the Martini et al cohort (**Figure 4E**). Of these contigs, four were partially anchored to GRCh38 reference, whereas six were unplaced.

### Sequence characteristics and functional landscape of below-threshold APG contigs

21,187 APG contigs failed to meet the NP alignment criteria when mapped to the T2T-CHM13 and 47 HPRC linear reference assemblies (**Figure 4A**). We therefore re-assessed their mappability using the more permissive RG alignment threshold. Under these criteria, 14,796 contigs (∼25 Mbp) aligned to both the T2T-CHM13 and HPRC assemblies, while 5,649 contigs (10.4 Mbp) aligned exclusively to HPRC assemblies. The remaining 742 contigs, totaling ∼1.5 Mbp, either failed to align or were aligned at below RG threshold in all references. These sequences were designated as “below-threshold contigs” (BTCs) (**Figure 5A, Table S14**).

**Figure 5.**
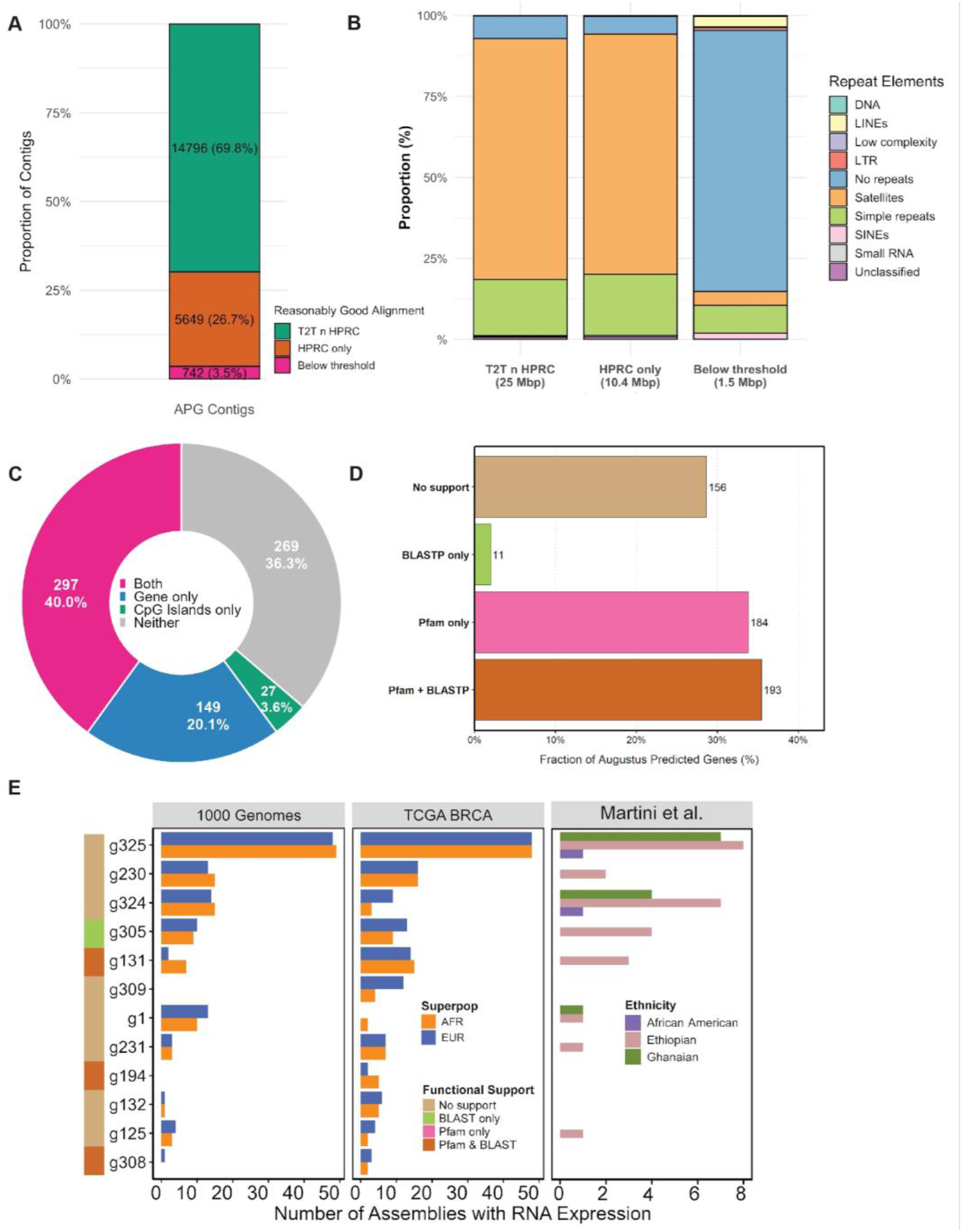
Functional characterization of APG contigs weakly or unmapped to gapless reference assemblies. (A) Mapping categories of APG contigs to T2T and HPRC at Reasonably Good Threshold. (B)Repeat profiles of APG contigs across mapping categories. (C) Proportions of functional feature classes for below-threshold contigs (BTC), which lack confident alignment to any reference genomes utilized. Category classes are shown in the legend, including genes and/or CpG islands (“Both,” “Gene only,” “CpG Islands only,” “Neither”). (D) Protein-level evidence of BTC predicted genes. Augustus gene models were assessed using Pfam domain annotation and BLASTP homology and categorized as supported by Pfam alone, BLASTP alone, by both, or unsupported. (E) Expression of BTC predicted genes across 3 diverse cohorts: 1000 Genomes Project, TCGA breast cancer, and Martini et al. The x-axis represents the number of samples per cohort expressing each of the displayed genes. Only the top 12 genes with the highest representation (TPM>0) across the 3 cohorts are shown. Functional prediction per gene is also provided as a row annotation.

To evaluate whether repeat content influenced this mappability pattern, we annotated repeat elements across all contig subsets using RepeatMasker (**Table S15**). Contigs mapping to both T2T-CHM13 and HPRC assemblies were highly enriched for repetitive sequences, with over 93% of their total length composed of repeats (**Figure 5B**). These sequences were primarily comprised of satellite (∼74%) and simple (∼17–19%) repeats, consistent with centromeric and subtelomeric genomic regions. In contrast, the 742 BTCs exhibited substantially lower repeat content, averaging ∼19%, with more than 80% lacking detectable repetitive elements (**Figure 5B, Table S15**). LINEs (3.2%), SINEs (1.9%), and LTR elements (0.89%) were the most prevalent repeats identified within BTCs.

Despite their limited mappability in linear reference space, BTCs showed substantial evidence of functional potential. Of the 742 contigs analyzed, 473 (63.7%) contained at least one functional feature, including predicted genes, CpG islands, or both **(Figure 5C, Table S16)**. Computational gene prediction identified coding potential in 446 contigs (60.1%), while CpG island detection revealed regulatory features in 324 contigs (43.7%). Notably, 297 contigs (40.0%) contained both predicted genes and CpG islands, suggesting coordinated coding and regulatory architecture rather than low-complexity or artifactual sequence. Only 269 contigs (36.3%) lacked detectable gene or CpG features. In total, these BTCs encoded 544 predicted gene models and 573 CpG islands spanning 344.8 kb, indicating that sequences poorly represented in linear references collectively harbor substantial predicted coding and regulatory content.

Protein-level validation further supported the biological relevance of predicted genes encoded within BTCs **(Figure 5D)**. Among predicted genes, 377 (70.8%) showed evidence of protein-coding potential based on Pfam domain annotation and/or BLASTP homology. Strong support from both Pfam and BLASTP was observed for 193 genes (36.2%), while 184 genes (34.6%) were supported by Pfam domains alone and 11 genes (2.1%) by BLASTP alone. Only 156 predicted genes (29.2%) lacked detectable protein support. The predominance of Pfam-supported predictions suggests that many encoded proteins belong to conserved domain families, even when full-length homologs are not readily identifiable by sequence similarity.

### RNA-seq mapping confirms that BTCs encode actively transcribed genes

To evaluate whether BTCs contain transcriptionally active sequences, we aligned RNAseq reads that failed to map to GRCh38 against a reference panel of BTC sequences using the predicted BTC genes as annotations. BTC predicted gene expression was detected in 98/100 1KGP RNAseq samples, with 18 of the 544 predicted BTC genes expressed across multiple samples. Predicted gene g325, detected in 97 samples, showed the most widespread expression (**Figure 5E**). Several expressed BTC genes exhibited BLAST and/or Pfam support for protein-coding potential, although *g325* lacked clear functional annotation despite its ubiquitous expression (**Figure 5E**).

To assess functional potential in a disease context we examined TCGA BRCA RNAseq dataset. BTC gene expression was detected in 99 of 100 samples across 36 predicted BTC genes, including 14 of the 18 genes observed in the 1000 Genomes samples. Expression patterns were concordant between cohorts (**Figure 5E**), with predicted genes *g305* and *g131* showing both widespread expression and protein-coding functional support. Analysis of the Martini et al. breast cancer cohort identified BTC expression in 26/43 samples, with 18 samples showing gene-level expression across 10 BTC genes, eight of which overlapped with the most commonly expressed genes in the 1000 Genomes and TCGA cohorts. g325 remained the most frequently expressed BTC gene (**Figure 5D**).

Together, these results demonstrate that BTCs contain transcriptionally active loci absent from current reference genomes, including the HPRC linear assemblies, highlighting persistent gaps in the human genome representation, particularly for ancestrally diverse populations.

## Discussion

Human reference genomes are foundational to genetic discovery, yet they are incomplete and underrepresent global genomic diversity, particularly African diversity. A central implication of this study is that completeness and representativeness are distinct. While T2T-CHM13 resolves sequence gaps and HPRC improves recovery of ancestry-enriched APG contigs, neither fully captures the functional breadth of human genomic diversity. Sequences present only in HPRC assemblies or remaining unresolved are particularly consequential, as they are largely invisible to standard pipelines for read alignment, annotation, and variant interpretation. Recent studies now make this point harder to dismiss as a purely technical concern. Inclusive graph-based references reduce reference bias^24^, ancestry-matched references alter scRNA-seq gene assignment^25^, diverse long-read transcriptomics reveals systematic ancestry bias in gene annotation^21,26^, and pangenome-based analyses improve sensitivity for disease-relevant structural variants^27,28^. In this context, missing reference sequence represents a mechanism by which biologically relevant variation is systemically obscured, particularly in populations historically underrepresented in reference resource^29^. This distinction is critical as African genomes harbor the greatest human genetic diversity, yet remain underrepresented in the reference frameworks that shape discovery, annotation, and interpretation.

By systematically characterizing African Pan-Genome contigs derived from GRCh38 unmapped reads^14^, we show that even advanced linear assemblies, including T2T-CHM13 and HPRC assemblies, incompletely capture ancestry-enriched genomic sequences. Alignment to T2T-CHM13 placed approximately 40% of contigs, largely within genomic regions absent from GRCh38 and enriched for centromeric satellites and repetitive elements, consistent with longstanding challenges in assembling heterochromatic DNA using short-read technologies^30,31^. However, ancestry-specific sequences remained incompletely represented, reinforcing the limitation of single linear references for structurally complex and polymorphic loci. Mapping to HPRC assemblies substantially increased placement rates and revealed ancestry-associated enrichment, particularly within African genomes, reflecting both demographic history and the greater genetic diversity^32^. Importantly, many contigs overlapped annotated genes, CpG islands, and pathways related to immune and cellular signaling, indicating that sequence absent from current references can harbor biologically relevant elements. A subset of contigs remained poorly mapped across all assemblies yet exhibited predicted coding features and transcriptional evidence, demonstrating that functional sequence can remain undetectable in alignment-based analyses. Such reference bias reduces sensitivity for variant detection and downstream analyses, disproportionately affecting underrepresented populations^14,33,34^.

Several limitations of our study should be considered. Functional annotation of BTCs relied on computational predictions and may overestimate functional potential, particularly sequences lacking homologs in existing databases^35^. Although RNA-seq data support the expression of a subset of predicted genes, experimental validation across tissues and development contexts remains necessary. Additionally, the APG contigs were derived from an admixed African American cohort, which may limit generalizability to continental African populations. Uneven representation within the Phase I HPRC panel may also influence observed ancestry-associated patterns.

Despite these limitations, our findings have important translational implications. Sequences absent from widely used reference genomes can contain coding and regulatory elements, indicating that clinically relevant variants within these regions are systematically overlooked^34^. Incorporating ancestry-enriched sequence into population-aware and graph-based reference frameworks will improve variant discovery, gene annotation, and transcriptomic profiling, ultimately enhancing the accuracy and equity of precision medicine initiatives. In conclusion, characterization of APG contigs reveals that substantial functional, ancestry-structured genomic sequence remains beyond current reference genomes. Addressing these gaps will be critical to reducing reference bias, improving variant interpretation, and ensuring that genomic discoveries benefit all global populations.

## Methods

### African Pan Genome Contigs Data Sourcing

We accessed the published African Pan Genome contigs sequences at GenBank with accession code PDBU01000000. All APG contig annotations were available via supplementary data from Sherman et al (2019)^14^. 125,715 APG contig sequences were published, however, 1475 were annotated as “dead” and were therefore excluded in all subsequent analyses (**Table S1**). In all analyses described, we proceeded with the remaining 124,240 APG contig sequences.

### Mapping APG contigs to GRCh38, T2T-CHM13 and HPRC linear references

We sought to determine if any of the APG contig sequences are represented in the T2T-CHM13 and the Phase I HPRC linear reference assemblies (n = 47). We utilized a similar approach as outlined by Sherman et al^14^, to map and assess the representation of APG contigs among unique reference genomes representative of African, South American, European, East and South Asian ancestries **(Figure S1)**. Specifically, we aligned the APG contigs to GRCh38.p10 as a benchmark to the Sherman et al analysis^14^, the T2T-CHM13 v2.0 reference genome^3^, and the Phase I HPRC haploid paternal linear assemblies v2^4^. Alignment to the various reference genomes and assemblies was performed using bwa-mem v0.7.17 with default parameters^36^. Alignments to individual locations within at least 300 bp were “chained” and analyzed as a single mapping output.

### Assessment of APG contig mapping to GRCh38, T2T-CHM13 and the HPRC linear references

To assess the quality of APG contig alignment to the various reference genomes and assemblies, we determined the coverage and identity of each alignment using thresholds set by Sherman et al^14^. Coverage is defined as the fraction of aligned length over the entire contig length. Identity is the fraction of correctly aligned bases over the aligned length. For individual alignments within 300 bp, we took the aligned length to be the sum of each alignment length, and measured identity by taking the weighted average of identities across each alignment, using the following formulae:

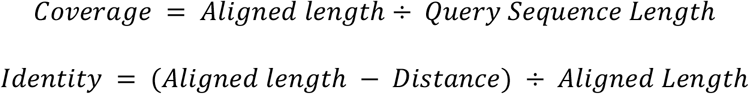

After calculating the coverage and identity, we applied two thresholds to determine if the mapping result was a “Reasonably Good” or “Nearly Perfect” alignment^14^. A Reasonably Good alignment was defined as having greater than or equal to 50% coverage, and greater than or equal to 80% identity.

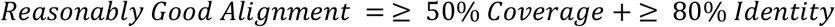

A Nearly Perfect alignment was defined as having greater than or equal to 80% coverage, and greater than or equal to 90% identity.

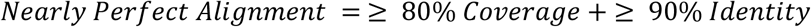

Using these metrics and thresholds, we determined the number of contigs passing each threshold for each reference assembly. A binary matrix was created to indicate the alignment status of each APG contig relative to T2T-CHM13 and each HPRC assembly, with values of 1 indicating alignment and 0 indicating no alignment/threshold for alignment not met.

To further assess the representativeness of APG contigs across HPRC superpopulations, the HPRC assemblies were categorized into superpopulation groups based on the designations from the 1000 Genomes Project: African (AFR), Admixed American (AMR), East Asian (EAS), European (EUR), and South Asian (SAS)^37^. For HPRC samples that were not a part of the 1000 Genomes project (HG002 and HG005), these were assigned to a corresponding superpopulation group (EUR and EAS, respectively) **(Table S9)**. A corresponding superpopulation-level and individual-level binary matrix was then generated, summarizing whether each contig aligned with at least one assembly within each superpopulation **(Table S12, S13)**. These matrices facilitated the evaluation of contig distribution and potential population-specific enrichment across HPRC assemblies. Additionally, to statistically assess the population-specific enrichment of APG contig alignments, a 2 × 5 contingency table was constructed, summarizing the counts of aligned and unaligned contigs across the superpopulations. The alignment counts were normalized according to the number of assemblies representing each superpopulation (AFR: 24, AMR: 16, EAS: 5, EUR: 1, SAS: 1) to account for the differences in sample sizes. A chi-square test of independence was conducted to determine the association between alignment status and population group. Odds ratios (ORs) were calculated for each pairwise comparison between AFR and other populations (AMR, EAS, EUR, SAS) to quantify the relative likelihood of contig alignment in AFR genomes.

### Functional enrichment of APG contigs across T2T-CHM13 genomic elements

To explore the potential localization of contigs mapped to T2T-CHM13 regions within functional genomic elements, we first obtained genomic coordinates for T2T-CHM13 unmasked CpG islands, NCBI RefSeq genes, problematic regions, as well as T2T-CMH13 unique regions in comparison to GRCh38 from the UCSC Genome Browser^38^ Table Browser (https://genome.ucsc.edu/cgi-bin/hgTables) in .*bed* file format. Additionally, we retrieved T2T-CHM13 RepeatMasker annotations generated by the T2T-CHM13 consortium (https://github.com/marbl/CHM13?tab=readme-ov-file). We then used BEDTools^39^ v2.30.0 to identify the presence of contigs across these various T2T-CHM13 genomic elements.

For co-localization of contigs with previously reported GWAS hits, we obtained GWAS associations from the EMBL-EBI GWAS Catalog (https://www.ebi.ac.uk/gwas/docs/file-downloads). GRCh38 GWAS variant positions were lifted over to obtain T2T-CHM13 coordinate system location by the T2T-CHM13 consortium (https://github.com/marbl/CHM13?tab=readme-ov-file). To identify genes associated with phenotypes or diseases among those mapped by contigs, we downloaded OMIM’s *Synopsis of the Human Gene Map* file (https://www.omim.org/downloads/) following licensing and registration, and extracted genes with reported phenotype or disease associations^19^.

### Gene ontology over-representation analysis

We extracted gene annotations using Ensembl gene models mapped to the T2T-CHM13 assembly, identifying a total of 373 genes that overlap with the regions where the APG contigs aligned. We utilized the R clusterProfiler package^40^ v4.14.6 to compare our test gene set against a background gene set composed of all annotated genes in the T2T-CHM13 assembly for enrichment. We evaluated the overrepresentation of Gene Ontology (GO) terms across functional categories, including Biological Process (BP), Cellular Components (CC), and Molecular Functions (MF). Statistical significance was assessed using Benjamini-Hochberg-adjusted p-values with a false discovery rate (FDR) of less than

0.05. We generated bar plots to summarize the enriched GO terms related to the genes within the APG-contig mapped regions.

### Assessment of Repetitive Elements in APG Contigs

Repetitive DNA elements in the African pan-genome (APG) contigs were identified using RepeatMasker^41^ v4.1.6, a widely used tool for detecting and classifying interspersed repeats and low-complexity sequences. The analysis was performed on three subsets of contig categories based on the Reasonably Good alignment threshold. These are: (i) those aligning to both T2T-CHM13 and HPRC assemblies, (ii) those aligning only to HPRC, and (iii) Below threshold contigs (those that failed to align or are aligned to either reference below the reasonably good threshold. RepeatMasker was executed using the Dfam 3.7 and RepBase repeat libraries^42,43^, utilizing the human-specific repeat database for precise classification. The analysis was performed with the parameters-species set to human to confine the search to repeats curated for the human genome.

### Gene and CpG Island Predictions and Annotation

We deployed AUGUSTUS v3.5.0^44^, an *ab initio* gene prediction software tool trained on human gene models to annotate potential protein-coding genes within BTCs. Contigs were individually processed, and gene structures including start and stop codons, exons, and coding sequences (CDS) were predicted. The resulting gene predictions were filtered to retain only transcripts with complete CDS features, representing putative protein-coding genes for downstream annotation (**Table S16**).

Additionally, to assess the epigenetic potential of APG contig sequences, we used CpGplot v6.6.0.0, a component of the European Molecular Biology Open Software Suite (EMBOSS, https://www.ebi.ac.uk/Tools/seqstats/emboss_cpgplot/). This tool detects CpG-rich sequences based on windowed GC content and the observed-to-expected CpG dinucleotide ratio. We used default thresholds commonly applied in genome-wide CpG prediction: a minimum island length of 200 bp, GC content ≥ 50%, an observed/expected CpG ratio of ≥ 0.6, and a 100 bp sliding window. The identified CpG islands were filtered to show only contigs with islands, where we quantified the number of islands predicted. The output was parsed and converted to BED format for comparative positional analysis and overlay with predicted genes (**Table S16**).

Translated protein sequences derived from AUGUSTUS gene models were queried against curated protein domain databases using profile hidden Markov models to identify conserved functional motifs and domain architectures, providing evidence for protein-coding plausibility beyond open reading frame structure alone^45^. In parallel, homology-based searches were performed against comprehensive protein sequence databases using sequence similarity alignment to detect significant matches to known proteins, allowing inference of evolutionary conservation and partial or divergent homology^46^.

### Quantification of Expressed Contig Sequences

To assess evidence of expressed sequences in APG contigs, we further analyzed RNA-seq data from three cohorts: (i) 1000 Genomes lymphoblastoid cell lines (n = 100; 52 African and 48 European)^37^; (ii) The Cancer Genome Atlas (TCGA) Breast Cancer (BRCA) primary tumors samples^22^ (n = 100; 50 AFR and 50 EUR) using a genetic admixture level > 0.85)^47^; and (iii) primary breast tumor samples an African ancestry enriched cohort reported in Martini et al^23^(n = 44; 9 African-American, 19 Ethiopian, 13 Ghanaian, 3 White). RNA-seq data was acquired in the following ways: 1000 Genomes .*fastq* files were compressed and aligned to GRCh38 genome version (*Homo_sapiens_assembly38*.*fasta* derived from the GATK resource bundle: https://console.cloud.google.com/storage/browser/genomics-public-data/resources/broad/hg38/v0;tab=objects?prefix=&forceOnObjectsSortingFiltering=false) using the *star_salmon* aligner and unmapped reads retrieved. For the TCGA BRCA samples, we obtained GRCh38 unmapped reads in a compressed .*fastq* format from the GRCh38 aligned .*bam* files we obtained from GDC Commons, using *samtools* v1.21 to retrieve only the unmapped reads.

To assess whether HPRC aligned contigs (n = 53,983) harbored expressed sequences, we used STAR v2.7.11b first to generate an index for the reference .*fasta* file containing the HPRC-mapped only contigs sequences, and then to align the RNA-seq data from all three cohorts to the same contigs. Reads per million mapped (RPM) normalized reads were obtained and stranded bedgraph output files were used for downstream analyses. To evaluate expression of predicted BTC genes, GRCh38-unmapped reads from the same cohorts were also aligned to a reference containing all BTCs. RNA-seq processing was performed using the *Nextflow* v25.10.0 nf-core/rnaseq pipeline v3.21.0. The AUGUSTUS-derived gene models provided in GTF format were used as gene annotations for the expression quantification step, yielding read counts and transcripts per million (TPM) values per predicted gene and sample. Because some samples were expected to lack detectable BTC expression, pipeline execution was configured to continue processing samples for which STAR reported zero aligned reads to the BTC reference genome.

## Acknowledgements

This work was made possible by the MacMillan Family Foundation as part of the MacMillan Center for the Study of the Non-Coding Cancer Genome at the New York Genome Center. This work was performed using the CPU and GPU compute nodes of the high-performance computing (HPC) cluster at Cold Spring Harbor Laboratory (CSHL). This cluster is supported by US National Institutes of Health grant S10OD028632-01, P01CA272295, R01CA266279.

We would like to acknowledge the Human Pangenome Reference Consortium (BioProject ID: PRJNA730823) and its funder, the National Human Genome Research Institute (NHGRI). Some of the results shown here are in part based upon data generated by the TCGA Research Network: https://www.cancer.gov/tcga. We are grateful to our colleagues for their thoughtful discussions and constructive feedback, which helped strengthen this study.

## Author Contributions

**Conceptualization**: RM, AT, KF, MBD, NC; **Methodology**: RM, DC, AT, KF, AA, MBD, NC; **Software, Formal analysis**: RM, AT, KF, DC, SM, AA; **Data Curation**: RM, AT, KF, DC, AA; **Visualization**: RM, AT, KF, DC, AA; **Writing - Original Draft**: RM, AT, KF, NC; **Writing - Review & Editing**: RM, AT, KF, DC, CEM, ICC, NR, SM, JW, OB, MBD, NC; **Supervision:** RM, NC, MBD; **Project administration:** RM, NC; **Funding acquisition**: RM, NC, OB, MBD

## References

1. Li, H. et al. Exome variant discrepancies due to reference-genome differences. Am. J. Hum. Genet. 108, 1239–1250 (2021).

2. Nie, J., Tellier, J., Tarasova, I., Nutt, S. L. & Smyth, G. K. T2T-CHM13 versus hg38: accurate identification of immunoglobulin isotypes from scRNA-seq requires a genome reference matched for ancestry. NAR Genom. Bioinform. 7, lqaf074 (2025).

3. Nurk, S. et al. The complete sequence of a human genome. Science 376, 44–53 (2022).

4. Liao, W.-W. et al. A draft human pangenome reference. Nature 617, 312–324 (2023).

5. Miga, K. H. & Wang, T. The need for a human pangenome reference sequence. Annu. Rev. Genomics Hum. Genet. 22, 81–102 (2021).

6. Clavell-Revelles, P. et al. Long-read transcriptomics of a diverse human cohort reveals widespread ancestry bias in gene annotation. Genomics (2025).

7. Rosenfeld, J. A., Mason, C. E. & Smith, T. M. Limitations of the human reference genome for personalized genomics. PLoS One 7, e40294 (2012).

8. Fatumo, S. et al. A roadmap to increase diversity in genomic studies. Nat. Med. 28, 243–250 (2022).

9. Popejoy, A. B. & Fullerton, S. M. Genomics is failing on diversity. Nature 538, 161–164 (2016).

10. Martin, A. R. et al. Human demographic history impacts genetic risk prediction across diverse populations. Am. J. Hum. Genet. 100, 635–649 (2017).

11. Carlson, C. S. et al. Generalization and dilution of association results from European GWAS in populations of non-European ancestry: the PAGE study. PLoS Biol. 11, e1001661 (2013).

12. Aganezov, S. et al. A complete reference genome improves analysis of human genetic variation. Science 376, eabl3533 (2022).

13. Aganezov, S. et al. A complete reference genome improves analysis of human genetic variation. Science 376, eabl3533 (2022).

14. Sherman, R. M. et al. Assembly of a pan-genome from deep sequencing of 910 humans of African descent. Nat Genet 51, 30–35 (2019).

15. Horton, R. et al. Variation analysis and gene annotation of eight MHC haplotypes: the MHC Haplotype Project. Immunogenetics 60, 1–18 (2008).

16. Meyer, D., C Aguiar, V. R., Bitarello, B. D., C Brandt, D. Y. & Nunes, K. A genomic perspective on HLA evolution. Immunogenetics 70, 5–27 (2018).

17. Robinson, J. et al. IPD-IMGT/HLA database. Nucleic Acids Res. 48, D948–D955 (2020).

18. Ko, J., Na, M., Kim, S., Lee, J.-R. & Kim, E. Interaction of the ERC family of RIM-binding proteins with the liprin-alpha family of multidomain proteins. J. Biol. Chem. 278, 42377–42385 (2003).

19. McKusick-Nathans Institute of Genetic Medicine, John Hopkins University (Baltimore, MD). Online Mendelian Inheritance in Man, OMIM. https://omim.org/.

20. Daya, M. et al. Association study in African-admixed populations across the Americas recapitulates asthma risk loci in non-African populations. Nat. Commun. 10, 880 (2019).

21. Taylor, D. J. et al. Sources of gene expression variation in a globally diverse human cohort. Nature 632, 122–130 (2024).

22. Cancer Genome Atlas Network. Comprehensive molecular portraits of human breast tumours. Nature 490, 61–70 (2012).

23. Martini, R. et al. African ancestry-associated gene expression profiles in triple-negative breast cancer underlie altered tumor biology and clinical outcome in women of African descent. Cancer Discov. 12, 2530–2551 (2022).

24. Lin, M.-J., Iyer, S., Chen, N.-C. & Langmead, B. Measuring, visualizing, and diagnosing reference bias with biastools. Genome Biol 25, 101 (2024).

25. Yao, J. & Gazal, S. Evaluating genetic ancestry inference from single-cell transcriptomic datasets. HGG Adv. 7, 100564 (2026).

26. Clavell-Revelles, P. et al. Long-read transcriptomics of a diverse human cohort reveals ancestrybias in gene annotation. Nat Commun 16, 10194 (2025).

27. Groza, C. et al. Pangenome graphs improve the analysis of structural variants in rare genetic diseases. Nat Commun 15, 657 (2024).

28. Schloissnig, S. et al. Structural variation in 1,019 diverse humans based on long-read sequencing. Nature 644, 442–452 (2025).

29. Nassir, N. et al. A draft UAE-based Arab pangenome reference. Nat. Commun. 16, (2025).

30. Miga, K. H. Centromere studies in the era of ‘telomere-to-telomere’ genomics. Exp. Cell Res. 394, 112127 (2020).

31. Altemose, N. et al. Complete genomic and epigenetic maps of human centromeres. Science 376, eabl4178 (2022).

32. Campbell, M. C. & Tishkoff, S. A. African genetic diversity: implications for human demographic history, modern human origins, and complex disease mapping. Annu. Rev. Genomics Hum. Genet. 9, 403–433 (2008).

33. Chen, N.-C., Solomon, B., Mun, T., Iyer, S. & Langmead, B. Reference flow: reducing reference bias using multiple population genomes. Genome Biol. 22, 8 (2021).

34. Sirugo, G., Williams, S. M. & Tishkoff, S. A. The missing diversity in human genetic studies. Cell 177, 26–31 (2019).

35. Yandell, M. & Ence, D. A beginner’s guide to eukaryotic genome annotation. Nat. Rev. Genet. 13, 329–342 (2012).

36. Li, H. Aligning sequence reads, clone sequences and assembly contigs with BWA-MEM. arXiv [q-bio.GN] (2013).

37. 1000 Genomes Project Consortium et al. A global reference for human genetic variation. Nature 526, 68–74 (2015).

38. Perez, G. et al. The UCSC Genome Browser database: 2025 update. Nucleic Acids Res 53, D1243–D1249 (2025).

39. Quinlan, A. R. & Hall, I. M. BEDTools: a flexible suite of utilities for comparing genomic features. Bioinformatics 26, 841–842 (2010).

40. Xu, S. et al. Using clusterProfiler to characterize multiomics data. Nat. Protoc. 19, 3292–3320 (2024).

41. Smit, A. F. A., Hubley, R. & Green, P. RepeatMasker Open-4.0. (2013-2015).

42. Hubley, R. et al. The Dfam database of repetitive DNA families. Nucleic Acids Res. 44, D81–9 (2016).

43. Jurka, J. et al. Repbase Update, a database of eukaryotic repetitive elements. Cytogenet. Genome Res. 110, 462–467 (2005).

44. Stanke, M., Steinkamp, R., Waack, S. & Morgenstern, B. AUGUSTUS: a web server for gene finding in eukaryotes. Nucleic Acids Res 32, W309–12 (2004).

45. Finn, R. D. et al. The Pfam protein families database. Nucleic Acids Res. 36, D281–8 (2008).

46. Altschul, S. F., Gish, W., Miller, W., Myers, E. W. & Lipman, D. J. Basic local alignment search tool. J. Mol. Biol. 215, 403–410 (1990).

47. Carrot-Zhang, J. et al. Comprehensive analysis of genetic ancestry and its molecular correlates in cancer. Cancer Cell 37, 639–654.e6 (2020).

