## Supplemental-Figures-S1-S8 for "African Pan Genome Contigs Expose Biologically Relevant Sequence Still Hidden from Human Reference Frameworks"

- **Supplemental Figure 1.** African Pan Genome Contig Alignment Computational Workflow
- **Supplemental Figure 2.** GRCh38 chromosome placements and T2T-CHM13 chromosome mapping results for Reasonably Good mapping APG Contigs.
- **Supplemental Figure 3.** Placement of nearly perfectly and reasonably good aligned contigs across T2T-CMH13 chromosomes.
- **Supplemental Figure 4.** Overlap of APG contigs with GWAS hits across chromosomes.
- **Supplemental Figure 5.** Superpopulation-Level Summary of APG Contig Alignment Patterns Across HPRC Assemblies
- **Supplemental Figure 6.** RNAseq alignment to HPRC-mapped only contigs.
- **Supplemental Figure 7.** APG Contigs with Differential RNA-seq Coverage Across Populations in the 1000 Genomes Cohort
- **Supplemental Figure 8.** APG Contigs with Differential RNA-seq Coverage Across Populations in the TCGA BRCA Cohort

### **Supplemental Figure 1.** African Pan Genome Contig Alignment Computational Workflow

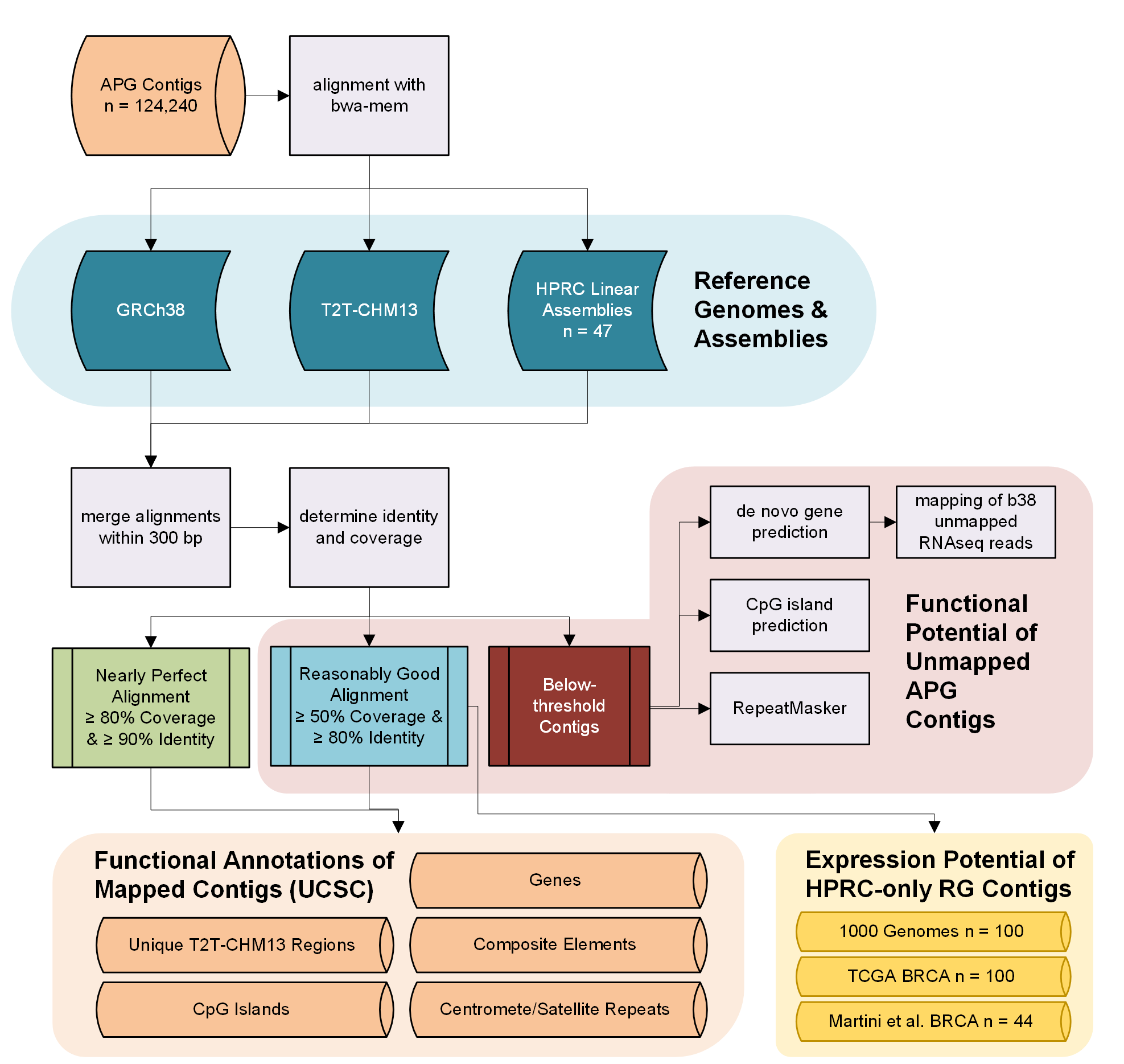

**Supplemental Figure 1.** African Pan Genome Contig Alignment Computational Workflow. Overview of the computational steps and resources utilized in the characterization of the 124,240 APG Contigs.

### **Supplemental Figure 2.** GRCh38 chromosome placements and T2T-CHM13 chromosome mapping results for Reasonably Good mapping APG Contigs.

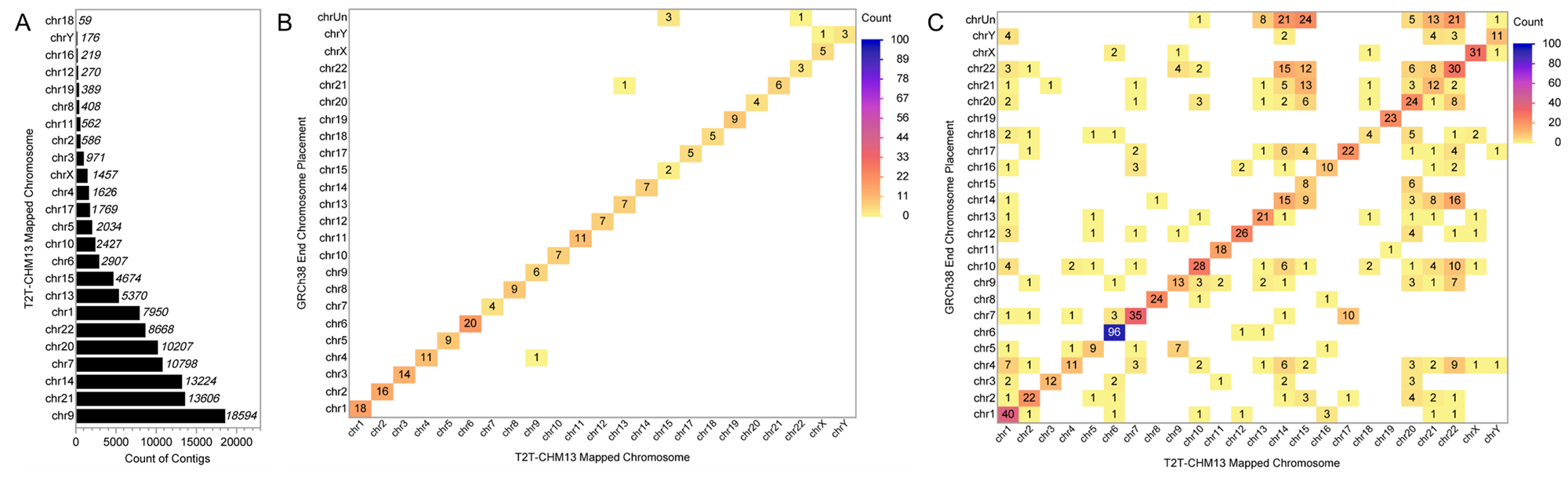

**Supplemental Figure 2.** GRCh38 chromosome placements and T2T-CHM13 chromosome mapping results for Reasonably Good mapping APG Contigs. (A) Bar chart of RG mapping to T2T-CHM13 chromosomes. Heatmaps of GRCh38 chromosome placements and T2T-CHM13 chromosome mapping results for (B) Two-End placed and (C) One-End placed APG contigs for Reasonably Good alignments.

### **Supplemental Figure 3.** Placement of nearly perfectly and reasonably good aligned contigs across T2T-CMH13 chromosomes.

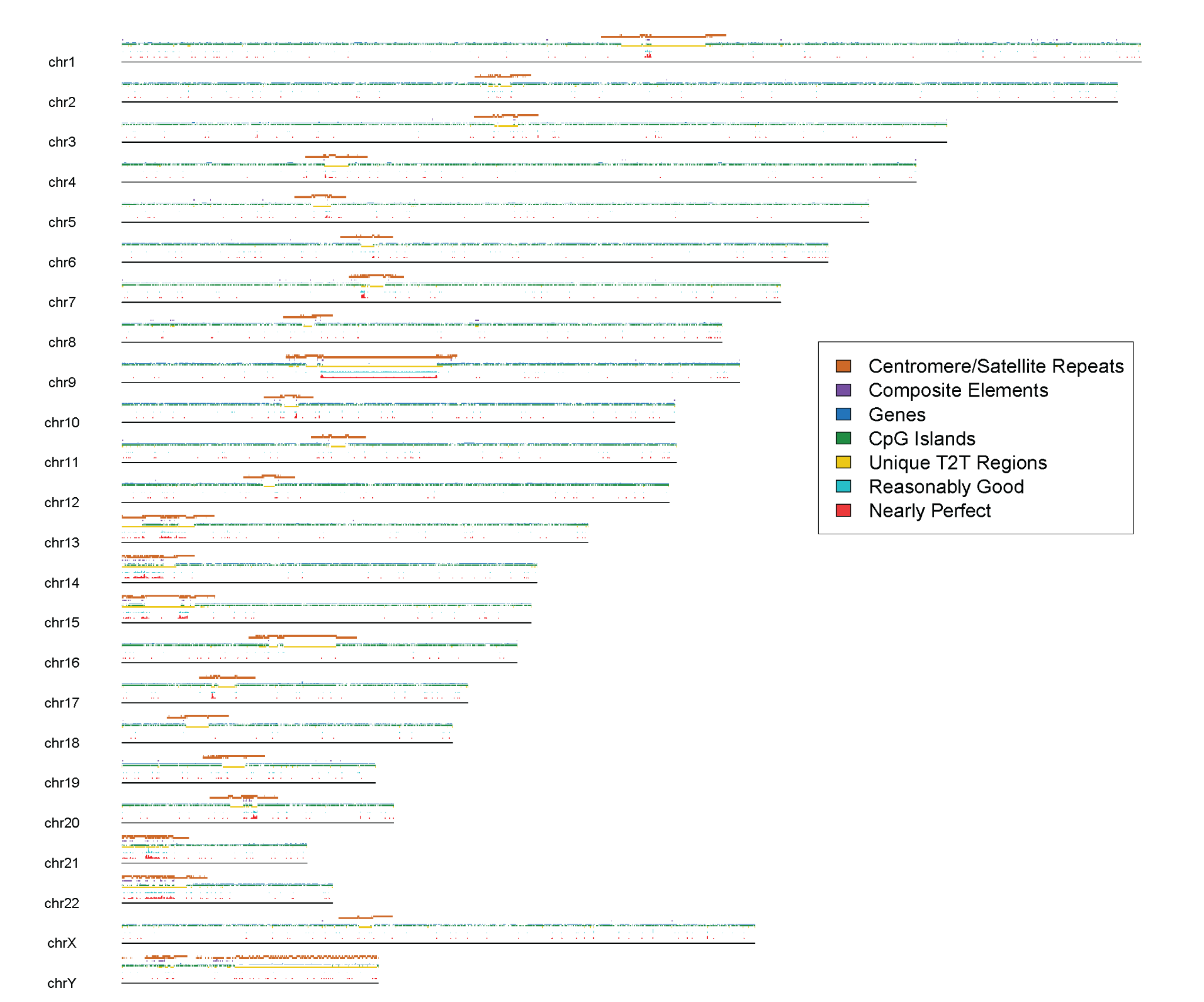

**Supplemental Figure 3.** Placement of nearly perfectly and reasonably good aligned contigs across T2T-CMH13 chromosomes. Nearly perfectly aligned contigs are highlighted in red, while reasonably good contigs are highlighted in turquoise. Enrichment was examined across T2T-CMH13 unique regions in comparison to GRCh38 (yellow), CpG Islands (green), RefSeq annotated genes (blue), composite elements (pink), and centromeres/satellite repeats (brown).

### **Supplemental Figure 4.** Overlap of APG contigs with GWAS hits across chromosomes.

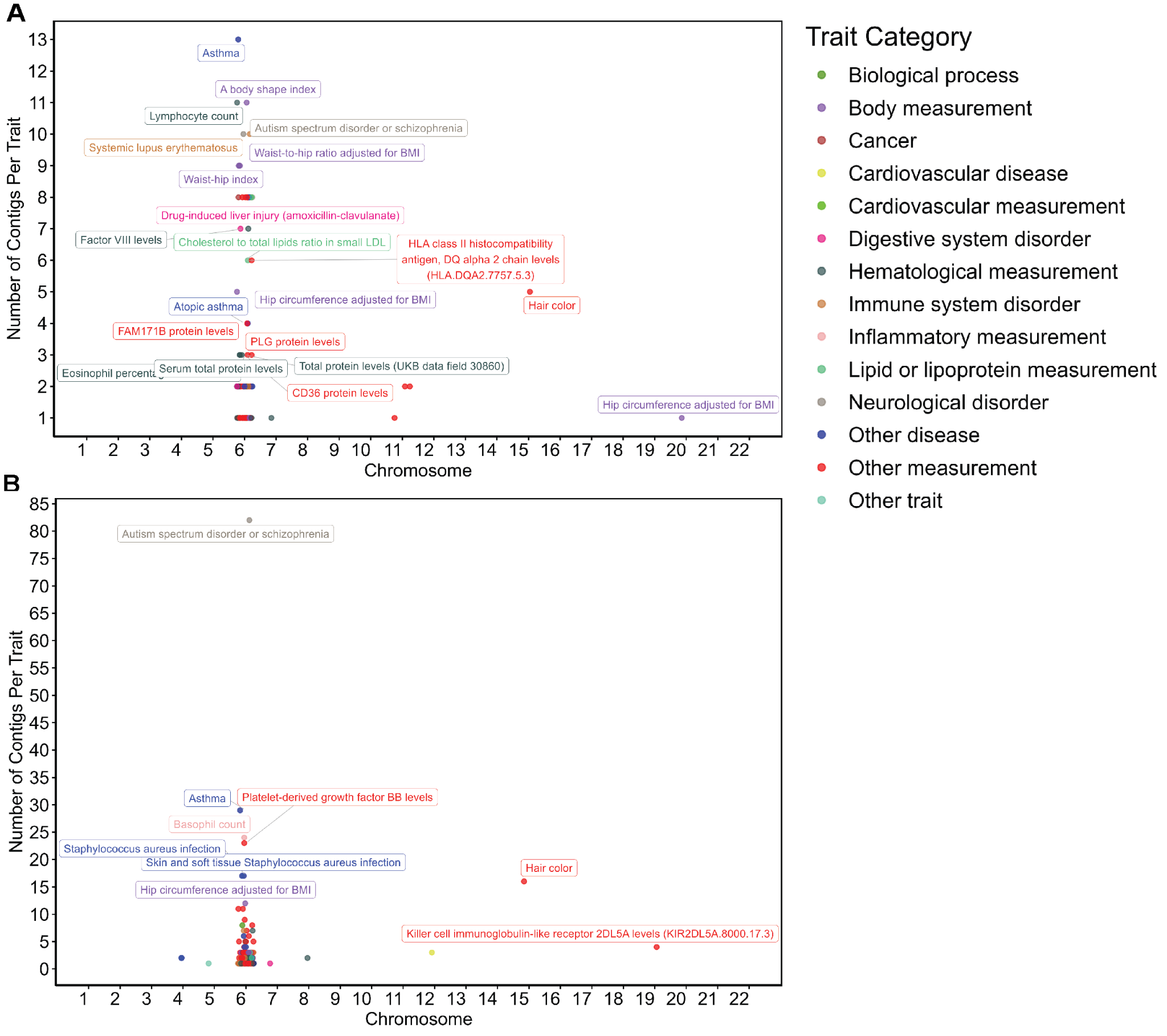

**Supplemental Figure 4.** Overlap of APG contigs with GWAS hits across chromosomes. Each dot represents a GWAS trait, with the y-axis indicating the number of contigs overlapping with GWAS hits (polymorphisms) for that trait. Trait labels are shown, and all traits are colored according to their parent trait category. Contig-GWAS hit overlap is visualized for (A) nearly perfect (NP), and (B) reasonably good (RG) alignments to T2T-CHM13.

### **Supplemental Figure 5.** Superpopulation-Level Summary of APG Contig Alignment Patterns Across HPRC Assemblies

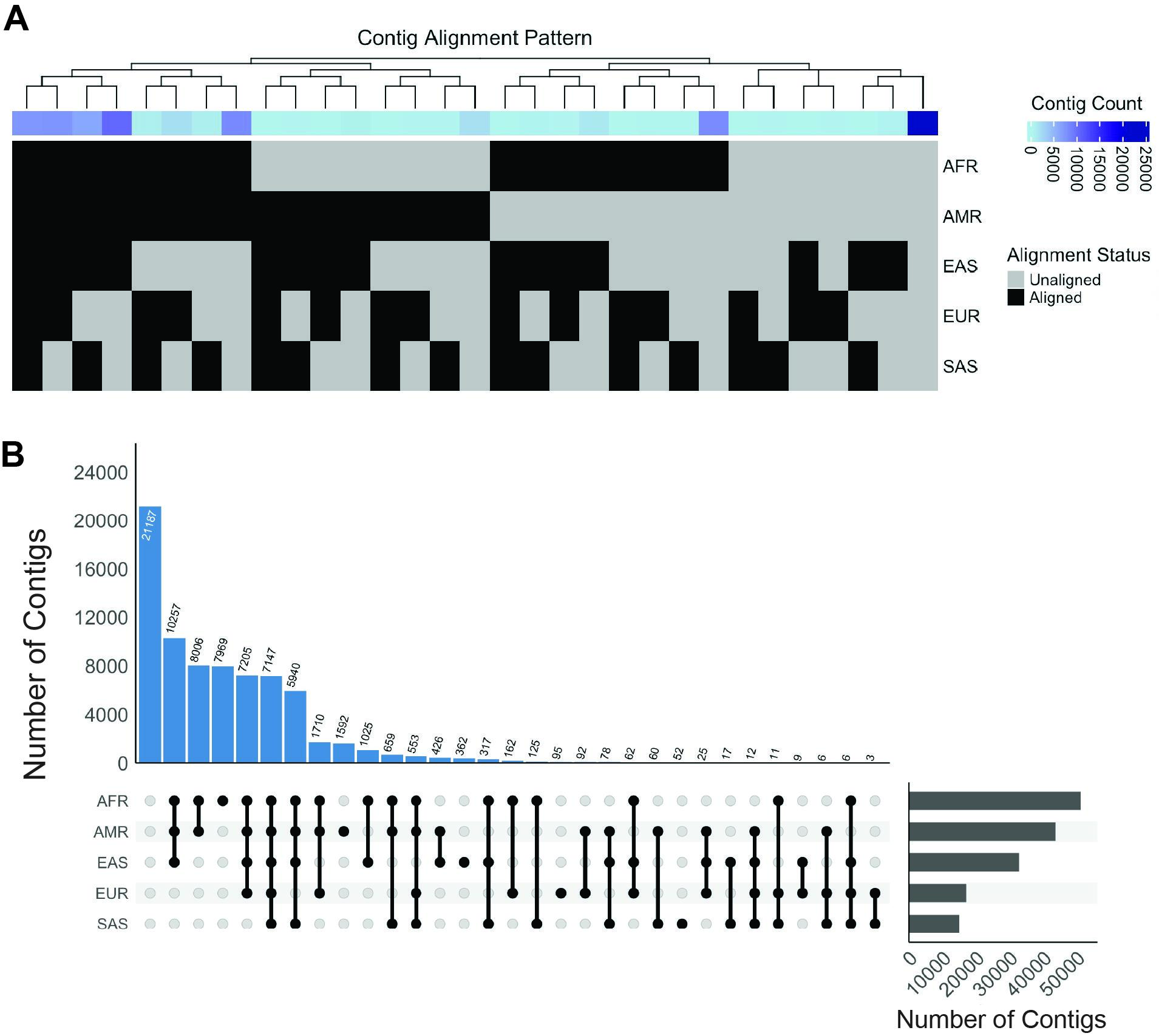

**Supplemental Figure 5.** Population-Level Summary of APG Contig Alignment Patterns Across HPRC Assemblies. (A) Continental-level binary alignment patterns across HPRC linear assemblies, where a contig is aligned to at least one member of each of the five human superpopulations. (C) Upset plot depicting the number of intersections of contig alignments across human superpopulations.

### **Supplemental Figure 6.** RNAseq alignment to HPRC-mapped only contigs.

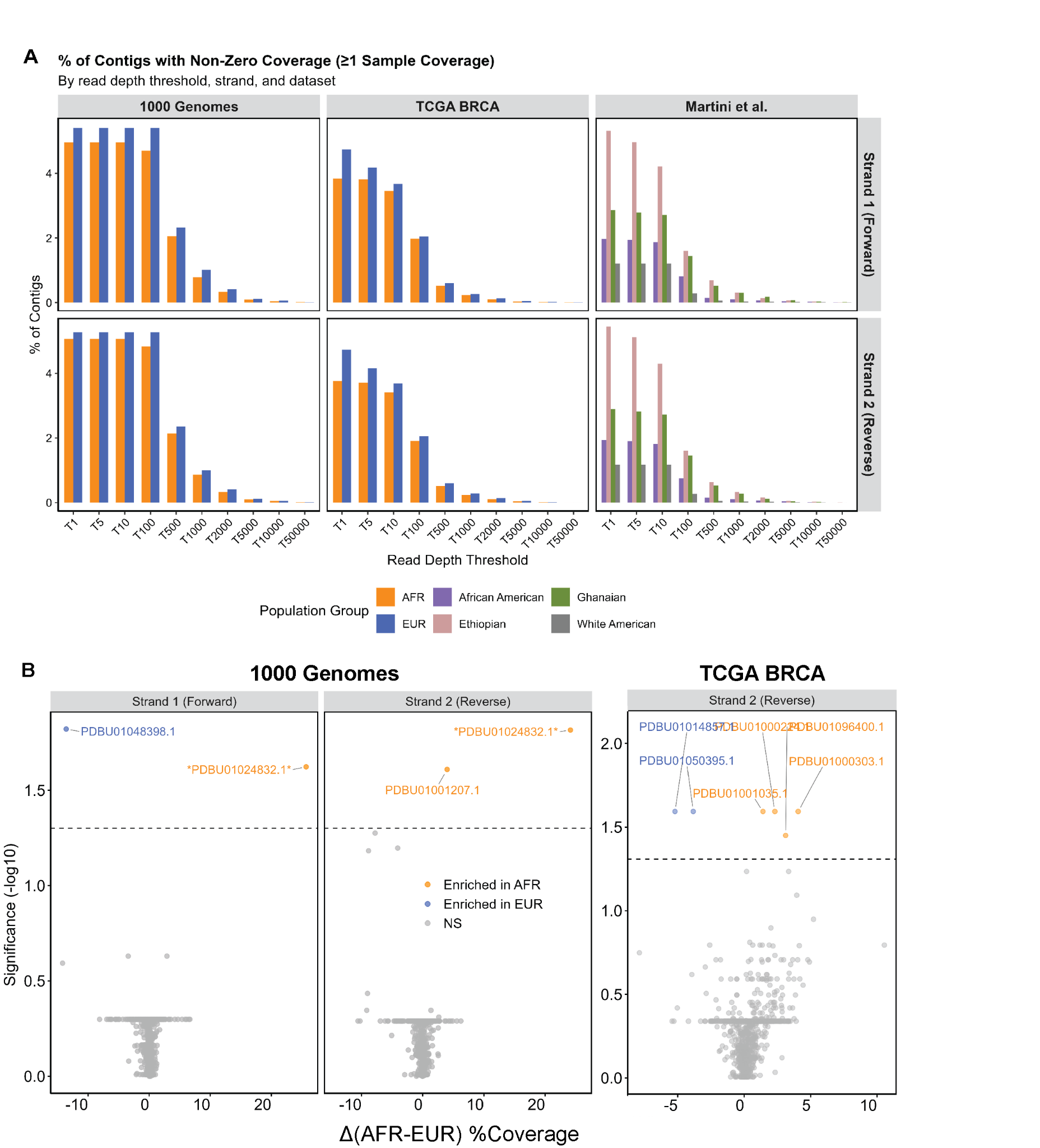

**Supplemental Figure 6.** RNAseq Alignment to HPRC-mapped only contigs. (A) Percentage of HPRC contigs (out of 53,983 total) exhibiting non-zero strand-specific coverage in at least one sample, stratified by read depth threshold (T1–T50,000), RNA strand (forward/reverse), and dataset. Three independent cohorts are shown: 1000 Genomes Project (AFR, n = 50; EUR, n = 50), TCGA breast cancer ( AFR, n = 52; EUR, n = 48), and Martini et al. (African American, n = 9; Ethiopian, n = 19; Ghanaian, n = 13; White American, n = 3), (B) Volcano plots showing differential strand-specific coverage between AFR and EUR populations at the T100 read depth threshold in the 1000 Genomes and TCGA BRCA, assessed by row-wise Wilcoxon rank-sum tests with Benjamini-Hochberg correction. The x-axis represents the difference in mean percent coverage (AFR − EUR), and the y-axis represents statistical significance (−log₁₀ adjusted p-value). Dashed horizontal line indicates padj = 0.05. Orange points denote contigs enriched in AFR; blue points denote contigs enriched in EUR; grey points are not significant (NS). Labeled contigs are those reaching significance (padj < 0.05); contigs significant on both strands are marked with asterisk.

### **Supplemental Figure 7.** APG Contigs with Differential RNA-seq Coverage Across Populations in the 1000 Genomes Cohort

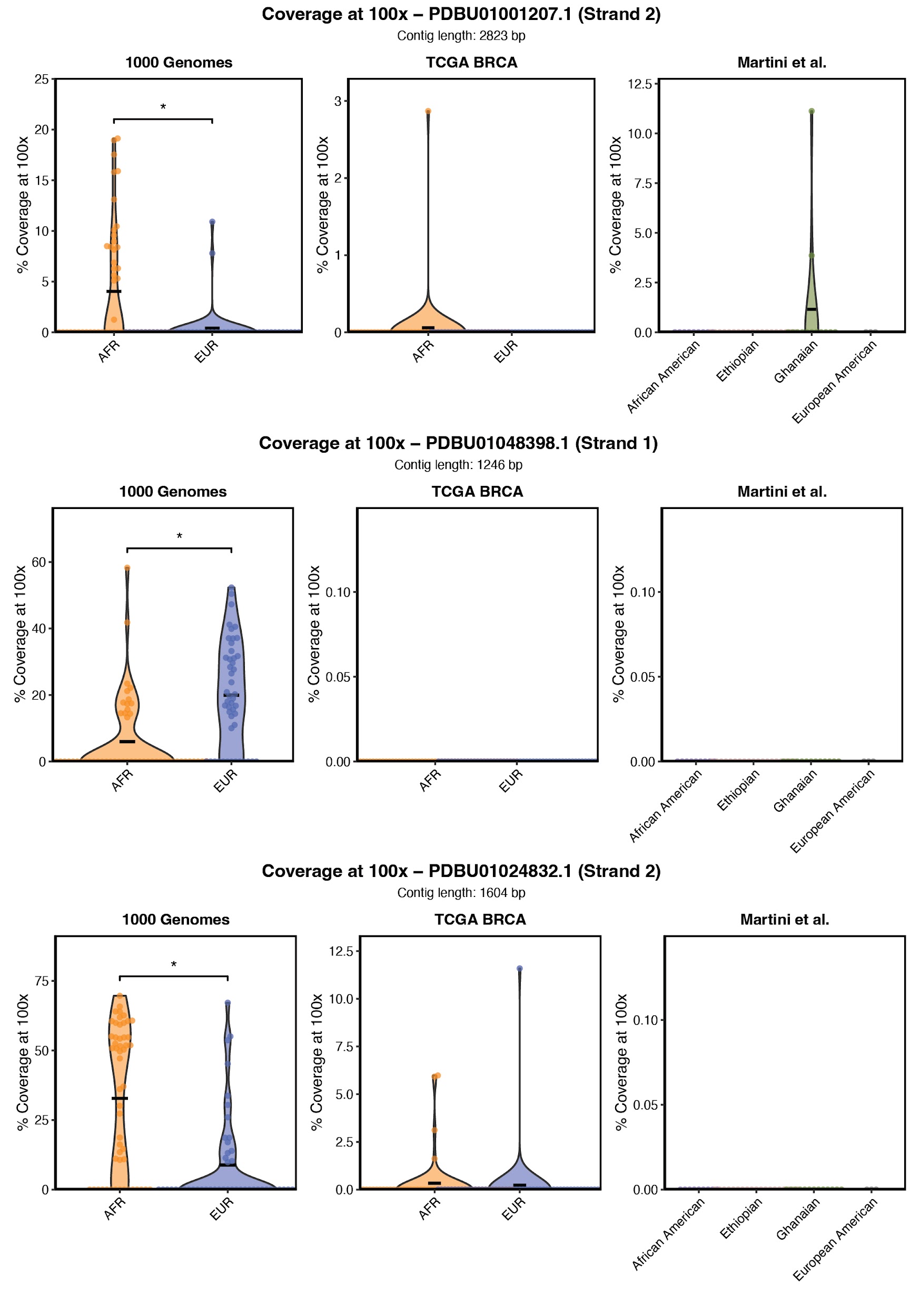

**Supplemental Figures 7.** APG Contigs with Differential RNA-seq Coverage Across Populations in the 1000 Genomes Cohort. Percentage of contig length covered at ≥100× RNA-seq coverage depth across all 3 studied cohorts for the remaining contigs identified as significantly differentially covered between AFR and EUR individuals not shown in Fig. 4E. Each point represents an individual sample, and samples are stratified by population group. Crossbar shows the mean of the distribution. Asterisks denote statistically significant differences between groups (*p* < 0.05 (*), *p* < 0.01 (**), or *p* < 0.001 (***)).

### **Supplemental Figure 8.** APG Contigs with Differential RNA-seq Coverage Across Populations in the TCGA BRCA Cohort

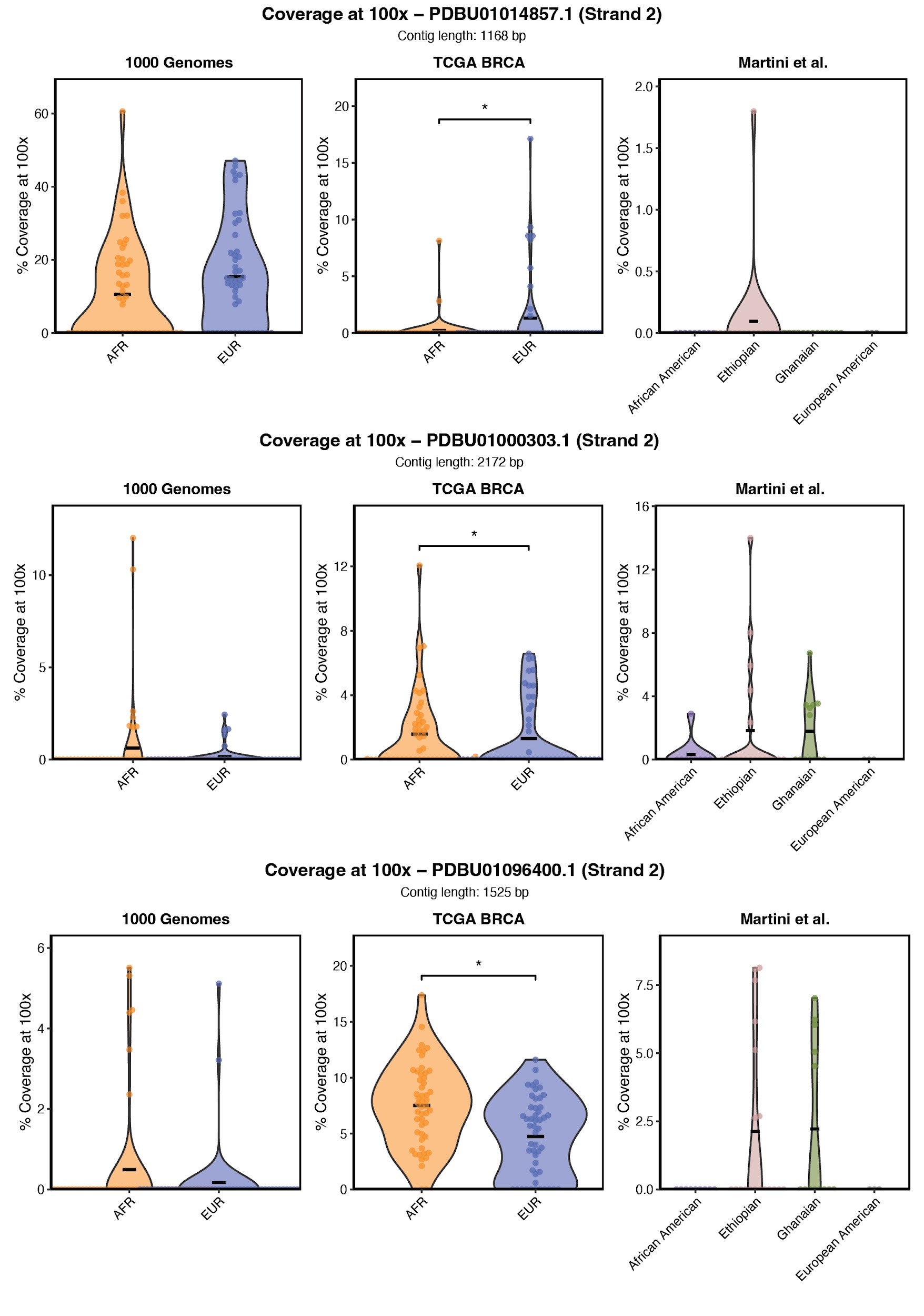

**Supplemental Figures 8.** APG Contigs with Differential RNA-seq Coverage Across Populations in the TCGA BRCA Cohort. Percentage of contig length covered at ≥100× RNA-seq coverage depth across all 3 studied cohorts for the remaining contigs identified as significantly differentially covered between AFR and EUR individuals not shown in Fig. 4E. Each point represents an individual sample, and samples are stratified by population group. Crossbar shows the mean of the distribution. Asterisks denote statistically significant differences between groups (*p* < 0.05 (*), *p* < 0.01 (**), or *p* < 0.001 (***)).
